# Functional study of the AMD-associated gene *TMEM97* in retinal pigmented epithelium using CRISPR interference

**DOI:** 10.1101/2020.07.10.198143

**Authors:** Jiang-Hui Wang, Daniel Urrutia-Cabrera, Santiago Mesa Mora, Tu Nguyen, Sandy Hung, Alex W. Hewitt, Thomas L. Edwards, Raymond C.B. Wong

**Affiliations:** Centre for Eye Research Australia, Royal Victorian Eye and Ear Hospital. Australia; Ophthalmology, Department of Surgery, University of Melbourne, Australia; Menzies Institute for Medical Research, School of Medicine, University of Tasmania, Australia

## Abstract

Age-related macular degeneration (AMD) is a blinding disease characterised by dysfunction of the retinal pigmented epithelium (RPE) which culminates in disruption or loss of the neurosensory retina. Genome-wide association studies have identified >65 genetic risk factors for AMD, including the *TMEM97* locus. *TMEM97* encodes the Sigma-2 receptor which is involved in apoptosis and cytotoxicity across a range of neurodegenerative diseases. However, the expression pattern of *TMEM97* in the human retina and its functional role in retinal cells has remained elusive. Here we utilised CRISPR interference (CRISPRi) to investigate the functional role of *TMEM97* in the retina. Transcriptome analysis of all major cell types within the human retina showed that *TMEM97* is expressed in the RPE, retinal ganglion cells (RGCs) and amacrine cells. Using CRISPRi, we performed loss-of-function study of *TMEM97* in the human RPE cell line, ARPE19. We generated a stable ARPE19 cell line expressing dCas9-KRAB which facilitated knockdown of *TMEM97* using specific sgRNAs. Our results show that knockdown of *TMEM97* in ARPE19 exerts a protective effect against oxidative stress-induced cell death. This work provides the first functional study of *TMEM97* in RPE and supports the role of *TMEM97* in AMD pathobiology. Our study highlights the potential for using CRISPRi to study AMD genetics, and the CRISPRi cell line generated here provided an useful *in vitro* tool for functional studies of other AMD-associated genes.

## Introduction

Age-related macular degeneration (AMD) is one of the leading causes of blindness in the western world with a prevalence of ~8.7% and projected to cause vision loss in 288 millions people worldwide by 2040 [1]. AMD is characterised by abnormalities of retinal pigment epithelium (RPE) at the macula region, leading to degeneration of the overlying photoreceptors and subsequently progressive loss of vision. Genetic and epidemiologic studies have greatly advanced our understanding of AMD pathology. Genome-wide association studies (GWAS) have successfully identified loci associated with genetic variants implicated with AMD risk and traits, with over 40 genetic variants being identified to be associated with AMD [2–6]. Of which, *APOE, CFH* and *ARMS2/HTRA1* are among the first identified loci to confer risk of developing AMD.

The *TMEM97/VTN* locus has been identified to infer significant risks of developing advanced AMD [2]. Five genes are present within the locus, including *TMEM97, VTN, POLDIP2, SLC13A2 and TMEM199*. In particular, *TMEM97* was recently identified to encode for the Sigma-2 receptor [7]. The Sigma-2 receptor plays a key role in mediating Ca+ and K+ signaling cascades in the endoplasmic reticulum and has been implicated in cancer development and a range of neurological degenerative diseases [8]. Activation of Sigma-2 receptor signaling has been shown to induce apoptosis and cytotoxicity [9]. Significant progress has been made to treat neurodegenerative diseases using Sigma-2 receptor inhibitors, including clinical trials in Alzheimer’s disease [10] and schizophrenia [11]. However, the expression pattern of *TMEM97* in different cell types within the human retina is not well studied. In addition, the functional role of *TMEM97* in the retina and its role in AMD remains unclear.

The emergence of CRISPR technology provides an efficient toolset for gene editing in the retina [12]. Moreover, recent development has highlighted the use of a nuclease-null Cas9 coupled with transcriptional repressors to modulate endogenous gene expression levels, termed CRISPR interference (CRISPRi), which offers a robust system to study gene function [13]. This study describes the use of CRISPRi for loss-of-function study for *TMEM97*. Using transcriptome analysis, we showed that *TMEM97* is expressed in multiple human retinal cell types, including RPE. We report the generation of a human RPE cell line, ARPE19, that stably expresses CRISPRi components (dCas9-KRAB). Using this CRISPRi system, we showed that knockdown of *TMEM97* exerts a protective effect on ARPE19 against oxidative stresses. Our results support a functional role of *TMEM97* in RPE degeneration, indicating that *TMEM97* could be a potential therapeutic target to treat AMD.

## Methods

### Donor retina collection

Collection of patient samples was approved by the Human Research Ethics committee of the Royal Victorian Eye and Ear Hospital (HREC13/1151H) and carried out in accordance with the approved guidelines. Post-mortem eye globes were collected by the Lions Eye Donation Service (Royal Victorian Eye and Ear Hospital) for donor cornea transplantation. The remaining eye globes were used to dissect the RPE/choroid layers for this study.

### Transcriptome analysis using RNAseq

Total RNA was extracted using the RNeasy kit and treated with DNase 1(Qiagen). Three donor samples of human RPE/choroid were processed for RNAseq. Transcriptome libraries and rRNA depletion were performed by the Australian Genome Research Facility. Samples were processed for 100bp single-end sequencing using an Illumina Novaseq 6000. Transcript level quantification was performed using *Salmon* to obtain gene-level counts [14].

### Cell culture

HEK293A cells were cultured in DMEM (Thermo Fisher, #11995-073), 10% [v/v] Fetal Bovine Serum (FBS), 1X GlutaMAX and 0.5% Penicillin/Streptomycin (all from Thermo Fisher). ARPE19 cells were cultured in DMEM/F12 (Gibco, #11320033;) supplemented with 10% FBS (Gibco, #10099141;) and 1% penicillin/streptomycin (Gibco, #15140122). All cells were grown at 37°C and 5% CO2 and passaged using 0.25% Trypsin-EDTA.

### Lentivirus generation

7×10^6^ HEK293FT cells were plated in a 10 cm dish one day prior transfection, and cultured with lentivirus packaging medium (Opti-MEM supplemented with 5% FBS and 200 μM Sodium pyruvate, all from Thermo Fisher). The lentivirus were generated by co-transfecting the dCas9-KRAB plasmid (gifts from Kristen Brennand; Addgene, #99372) with the three 3^rd^ generation packaging vectors pMDLg/pRRE (Addgene, #12251), pRSV-Rev (Addgene, #12253) pMD2.G (Addgene, #12259), using Lipofectamine 3000 (Thermo Fisher). The medium with lipofectamine 3000 was replaced with fresh media 6 hours after transfection. Afterwards, the crude virus was collected at 48 and 72 hours after transfection. The collected virus was filtered (0.45 μm filter, Sartorius) and concentrated using PEG-it overnight at 4°C (SBI Integrated Sciences) and resuspended in cold PBS.

### Puromycin dosage study

To determine the minimum concentration of puromycin required to select for ARPE19 cells transduced with lentiviruses, we treated ARPE19 cells with increasing concentrations of puromycin (250ng/ml, 500ng/ml, 1000ng/ml, 2000ng/ml). After 48 hours of treatment, we determined the lowest concentration of puromycin that was able to kill all ARPE19 cells and this dosage and duration was used for selection of the ARPE19-KRAB cell line.

### CRISPR interference (CRISPRi)

SpCas9 sgRNA expression cassettes containing a U6 promoter, sgRNA and sgRNA scaffold are synthesized, amplified and purified as previously described [15]. On Day 0, 6 x 10^4^ HEK293A cells or 1 x 10^5^ ARPE19-KRAB cells were plated down in a well on a 12-well plate. On day 1, the cells were transfected with 360ng sgRNA expression cassette and 800ng dCas9-KRAB, using Lipofectamine 3000 for HEK293A or RNAimax for ARPE19-KRAB. Mock control (no DNA transfected) was utilised as a negative control in every individual experiment. On day 4, the samples were processed for RNA extraction for qPCR analysis to assess gene expression levels.

### qPCR analysis

Total RNA were extracted using the Illustra RNAspin kit and treated with DNase 1 (GE Healthcare). cDNA synthesis and taqman qPCR were performed as previously described [16], using the following taqman probes for TMEM97 (Hs00299877_m1) and the housekeeping gene β-actin (Hs99999903_m1). The taqman assay was performed in a ABI 7500 or StepOne plus (Thermo Fisher).

### Immunocytochemistry

Standard immunocytochemistry procedures were carried out as we previously described [17]. Briefly, samples were fixed in 4% paraformaldehyde, followed by blocking with 10% serum (Sigma) and permeabilization with 0.1% Triton X-100 (Sigma). The samples were then immunostained with antibodies against Cas9 (Cell Signaling, #14697), Alexa Fluor 488 secondary antibodies (Abcam), and nuclear counterstain with DAPI (Sigma). Samples were imaged using a Zeiss Axio Vert.A1 fluorescent microscope.

### Cell viability assay

Cell viability assay was performed using a CellTiter-Glo^®^ Luminescent Cell Viability Assay (Promega, G9241;) following the manufacturer’s instructions. Briefly, 4×10^4^ cellsl were seeded in a well of a 96-well plate on Day 0, followed by CRISPRi on Day 1. For some conditions, the cells were treated with 100μM tert-Butyl Hydroperoxide (tBHP) on Day 3. On Day 4, old media was replaced with 25μl fresh media in each well, followed by adding 25μl CellTiter-Glo^®^ Reagent before incubating at room temperature for 10 minutes to initiate a luminescent reaction. 40μl of cell lysate from each well was loaded to an opaque white luminometer plate to measure luminescence using a Spark 20M microplate reader (Tecan).

### Short tandem repeat analysis

Genomic DNA of ARPE19 and ARPE19-KRAB were extracted using the Wizard SV Genomic DNA Purification kit (Promega), following manufacturer’s instructions. Short tandem repeat analysis is performed using the GenePrint 10 system (Promega) by the Australian Genome Research Facility.

### Statistical analysis

Statistical analysis is performed on biological repeats using paired t-test or one-way ANOVA with Tukey’s test (Graphpad Prism). p < 0.05 was used to establish statistical significance.

## Results

### Gene expression of TMEM97/VTN locus in human retina

First we set out to study the expression of genes in the TMEM97/VTN locus in the human retina. We performed *in silico* analysis using single cell RNA-seq data we previously reported for adult human neural retina [18]. Our results revealed that the retinal ganglion cells have strong expression of *TMEM97, POLDIP2* and to a lesser degree of *TMEM199* (Figure 1A). Interestingly, *POLDIP2* was also detected in low levels in a number of cell types, including amacrine cells and bipolar cells, Müller glia and cone photoreceptors. On the other hand, *VTN* and *SLC13A2* were mainly expressed in the cone photoreceptors (Figure 1A).

**Figure 1:**
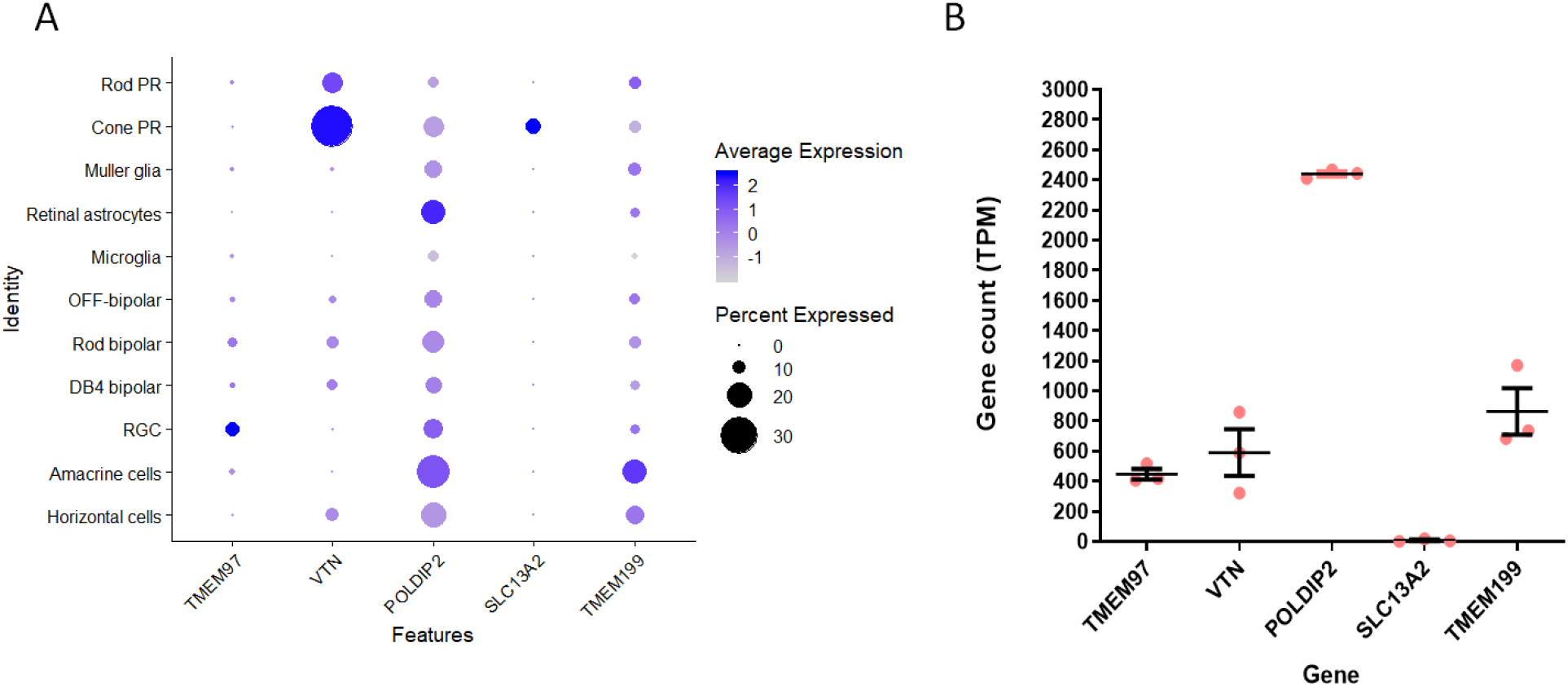
Expression of AMD-associated genes in human retina. Expression levels of AMD-associated genes in the *TMEM97/VTN* locus in A) human neural retina by single cell-RNAseq (n=3 donors;[18]). PR: photoreceptors; MG: Müller glia; RGC; retinal ganglion cells; and B) human RPE/choroid by RNAseq (n = 3 donors, error bars represent SEM).

To further determine the expression of TMEM97/VTN locus genes in RPE, we performed RNAseq in 3 human RPE samples (Figure 1B). The donor information is listed in Supplementary table 1. To avoid potential post-mortem effects, we selected donor RPE samples with a short retrieval time (4-6 hours). Our results showed that RPE express detectable levels of *TMEM97* (447±35 TPM)*, VTN* (590 ± 155 TPM)*, POLDIP2* (2440 ± 16 TPM) and *TMEM199* (863 ± 154 TPM). However, *SLC13A2* expression is extremely low in RPE (7 ± 5 TPM). Together, these results showed that most of the genes in *TMEM97/VTN* locus are expressed in both human neural retina and RPE. Next, we selected *TMEM97* as a candidate to perform further loss-of-function study in a human RPE cell line, ARPE19.

### Using CRISPRi to knockdown expression of *TMEM97*

To induce knockdown of *TMEM97* expression, we utilised the dCas9-KRAB system [19]. We designed 3 sgRNAs that target the proximity of the transcription start site (TSS) of *TMEM97* gene (Figure 2A). However, different databases (e.g. Refseq, UCSC, GENCODE) could annotate the TSS in different locations. A previous study by Radzisheuskaya *et al*. showed that sgRNAs designed using TSS annotated by the FANTOM/CAGE promoter dataset are the most reliable compared to other databases [20]. To test the positional effect of sgRNA target in CRISPRi, we designed a sgRNA using TSS defined by Ensembl (sgRNA 1), and 2 sgRNAs using TSS defined by the FANTOM/CAGE promoter dataset (sgRNA 2 and 3; Supplementary table 2).

**Figure 2:**
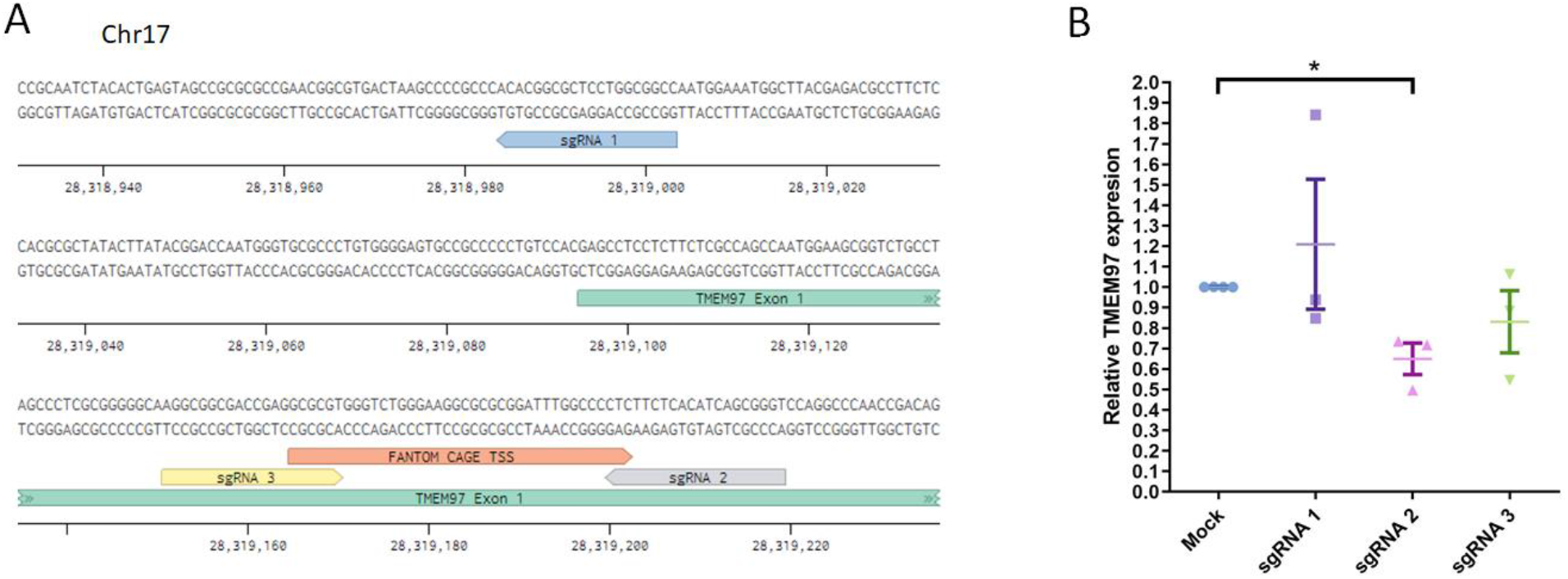
CRISPRi knockdown of TMEM97 expression. A) Schematic diagram of sgRNA target areas near the transcription start site (TSS) of human *TMEM97* gene. Green annotation marks the TSS and exon 1 for TMEM97 as determined by Ensembl. Orange annotation represents the TSS as determined by FANTOM CAGE data. B) qPCR analysis of CRISPRi knockdown of TMEM97 expression in HEK293 using 3 different sgRNA. Results represented as the mean ± SEM of three biological repeats, each with three technical repeats. * indicates p<0.05.

To test the sgRNA, we utilised an sgRNA expression cassette system synthesized as short 500-bp oligonucleotides [15]. We first evaluated the effect of the 3 sgRNA in modulating *TMEM97* expression in HEK293A cells. .qPCR analysis showed that sgRNA 1 did not induce knockdown of TMEM97 levels, while sgRNA 2 and 3 induced ~35% and ~15% knockdown respectively (Figure 2B, n=3). In addition, our results also showed that various doses of sgRNA did not have a significant effect on the knockdown levels of *TMEM97* (Supplementary figure 1). Thus, we utilised sgRNA 2 for further experiments to knockdown *TMEM97* expression.

### Generation of a stable ARPE19 cell line for CRISPRi

To facilitate the use of CRISPRi in retinal research, we generated a human RPE cell line that stably expresses dCas9-KRAB. ARPE19 cells were transduced with lentiviruses that constitutively express dCas9-KRAB under the EF1α promoter. The transduced ARPE19 cells were selected using 2ug/ml puromycin, as determined by an puromycin dosage study (Supplementary Figure 2). Following puromycin selection, the transduced ARPE19 cells (ARPE19-KRAB) were further validated for expression of dCas9-KRAB. Our results showed that ARPE19-KRAB expressed the KRAB component using RT-PCR analysis (Figure 3A). Furthermore, immunocytochemistry analysis showed that ARPE19-KRAB expressed detectable levels of SpCas9 protein (Figure 3B). Also, we performed short tandem repeat analysis and confirmed that ARPE19-KRAB originated from the parental ARPE19 cell line (Figure 3C). Together these results confirmed the successful generation of ARPE19-KRAB.

**Figure 3:**
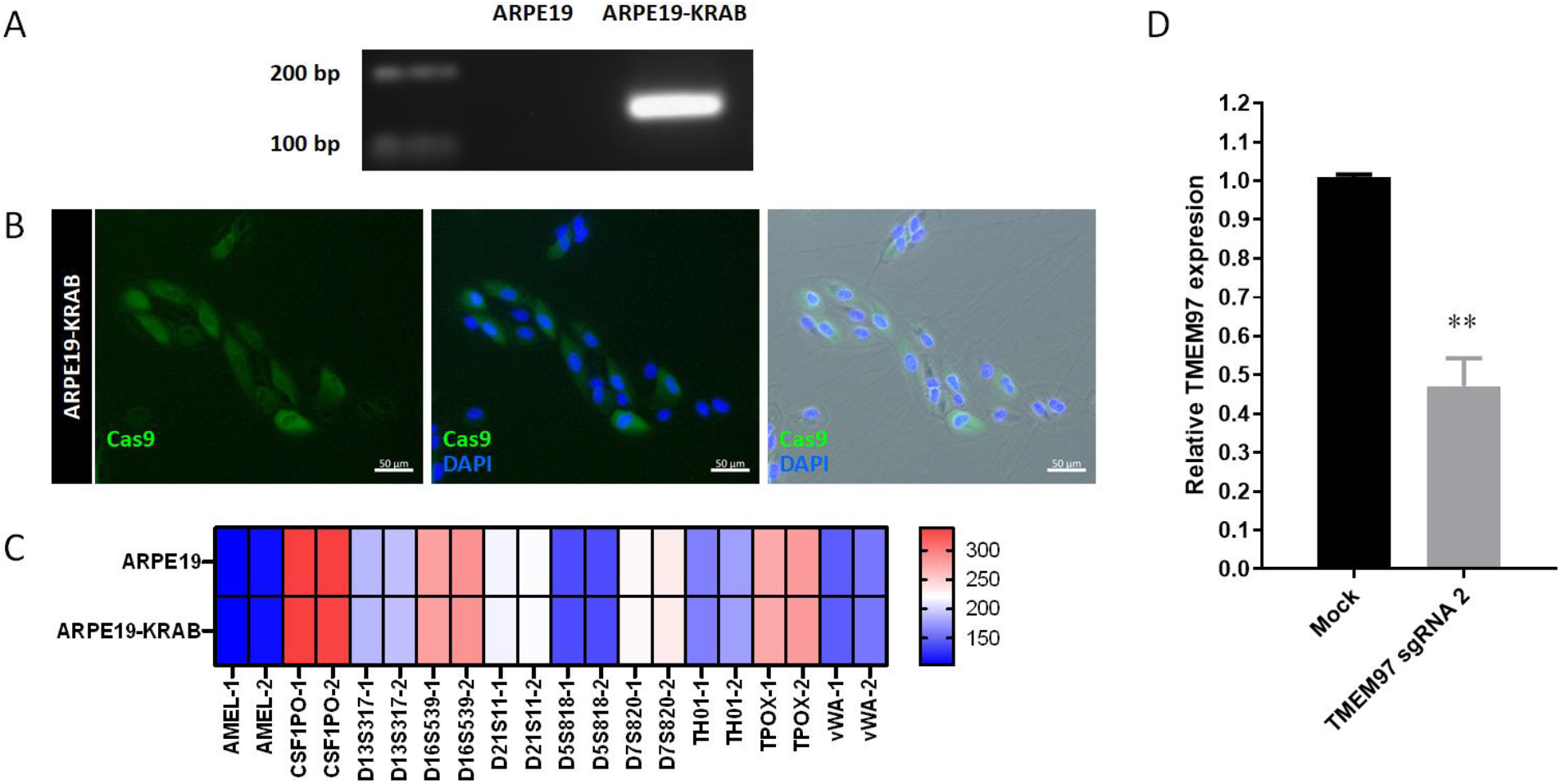
Generation of a stable CRISPRi human RPE cell line expressing dCas9-KRAB. Characterization of ARPE19-KRAB cell line showing expression of A) KRAB by RT-PCR, and B) SpCas9 by immunocytochemistry. C) Heatmap of short tandem repeat analysis of 10 polymorphic markers. Allele 1 and 2 are designated as −1 and −2 respectively. D) qPCR analysis of TMEM97 knockdown using CRISPRi in ARPE19-KRAB. Results are presented as the mean ± SEM of three biological repeats, each with three technical repeats. ** indicates p <0.01.

Next, we transfected sgRNA into ARPE19-KRAB using RNAimax to knockdown *TMEM97* expression. qPCR analysis showed that we could consistently achieve ~53% knockdown of *TMEM97* in ARPE19-KRAB using CRISPRi (Figure 3D, n=3). Notably, we observed that CRISPRi using a stable ARPE19 cell line expressing KRAB, achieved better knockdown efficiency compared to transfection in HEK293 cells. We have also tested transfection using Fugene HD, which showed a similar efficiency of *TMEM97* knockdown in ARPE19-KRAB (data not shown). These results showed that we have generated a robust *in vitro* system to perform loss-of-function study for *TMEM97* in human RPE cells.

### Loss of function study of TMEM97 in ARPE19 cells using CRISPRi

Using the ARPE19-KRAB system, we assessed if loss of *TMEM97* would affect cell viability of ARPE19. Our results demonstrated that CRISPRi-mediated knockdown of *TMEM97* did not affect cell viability of ARPE19 cells (Figure 4), suggesting that *TMEM97* might not be critical in maintaining RPE cell viability under normal conditions. Given that oxidative stresses play a key role in RPE degeneration in AMD [21], we further treated ARPE19-KRAB with tert-Butyl hydroperoxide (tBHP) as an oxidative stress to better mimic RPE under AMD conditions. Cell viability assay showed that tBHP treatment resulted in dramatic cell death in ARPE19-KRAB (0.33 ± 0.02 fold compared to control, Figure 4). Interestingly, *TMEM97* knockdown resulted in an significant increase of resistance to tBHP-induced cell death (0.66 ± 0.097 fold compared to control). Together, these results showed a novel role of *TMEM97* in regulating the viability of RPE and demonstrated the potential of using CRISPRi for functional studies of AMD-associated genes in RPE cells.

**Figure 4:**
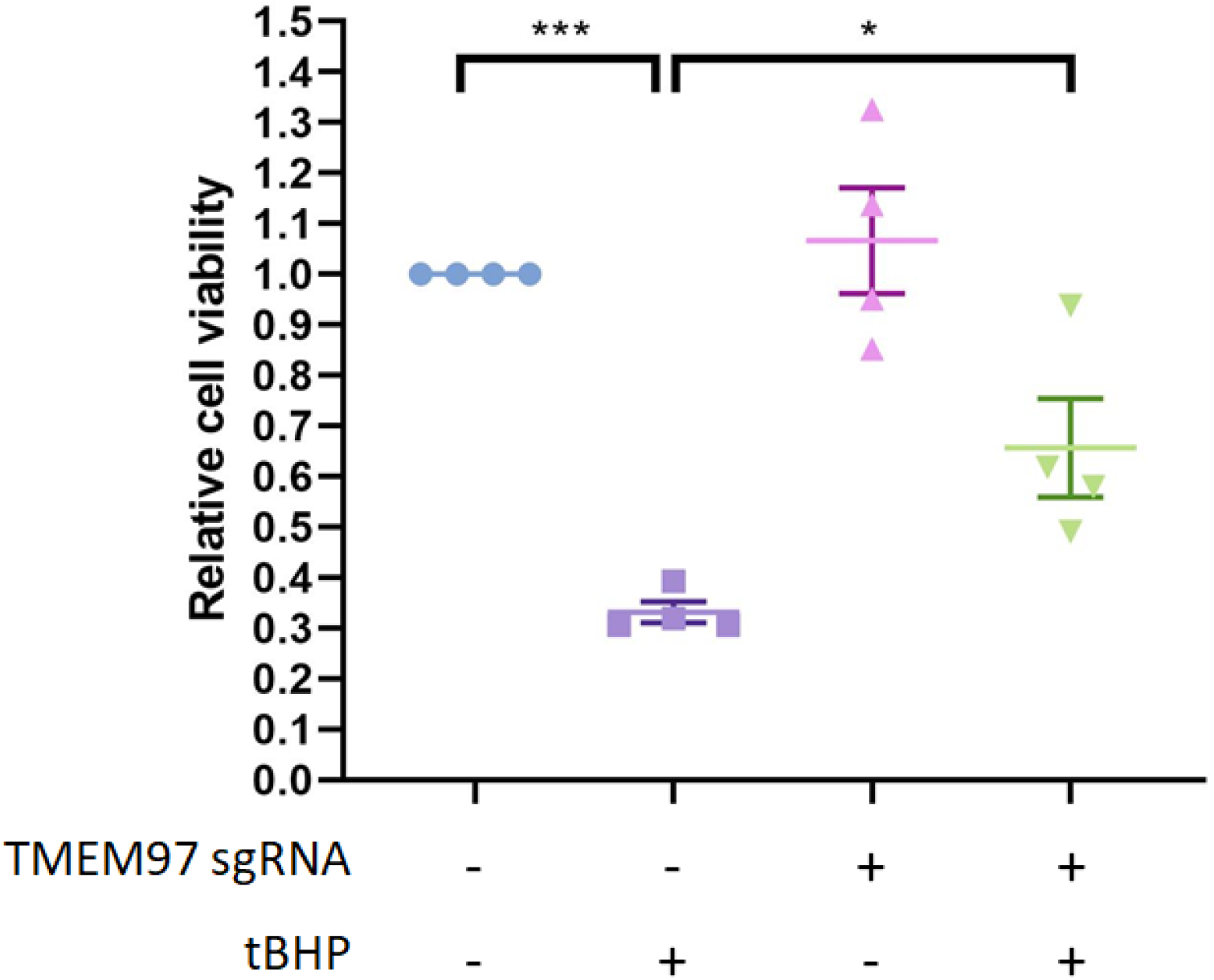
*TMEM97* knockdown exerts a protective effect on ARPE19 against oxidative stresses. Cell viability analysis of ARPE19-KRAB following *TMEM97* knockdown in the presence or absence of tBHP. Results are presented as the mean ± SEM of four biological repeats, each with four to six technical repeats. * indicates p<0.05; *** indicates p<0.001.

## Discussion

Advances in next generation sequencing have enabled the identification of genes associated with AMD, including the *TMEM97* loci [2]. However, we lack detailed knowledge of how AMD-associated genes impact specific retinal cell types in humans. In particular, the expression pattern and the function of *TMEM97* in specific retinal cells in humans remains unclear. Using transcriptome analysis of healthy human retina, our study showed that *TMEM97* is expressed in human RPE/choroid, as well as the RGC. Our data also revealed that other genes within the *TMEM97* loci are expressed in the RPE or the photoreceptors: *VTN* is expressed in both photoreceptors and RPE/choroid, *SLC13A2* is expressed in photoreceptors, *POLDIP2* and *TMEM199* are mainly expressed in RPE/choroid. These results support that the *TMEM97* loci genes may contribute to AMD development by acting on different retinal cell types.

Functional validation of putative causal genes would foster a better understanding of AMD etiology mechanisms, which is important to develop better diagnosis and treatment. However, study of AMD-associated genes in the retina of model organisms is challenging: the lack of macula in rodents represents a major limitation in modeling AMD, while primate studies are limited by the high research cost and the requirement of specialised housing facilities [22]. In this study, we present an *in vitro* human RPE model with CRISPRi to knockdown expression of *TMEM97*. We were able to identify sgRNA for successful knockdown using gene annotation from the FANTOM/CAGE study, which is consistent with a previous study [20]. Using this, we were able to identify a sgRNA that targeted the immediate downstream region of the TSS to knockdown expression of *TMEM97*. Moreover, we generated an ARPE19 cell line with constitutive expression of dCas9-KRAB, which can be combined with a sgRNA expression cassette system [15] to enable CRISPRi-mediated knockdown of gene expression. This provides an efficient tool to perform loss-of-function study for AMD-associated genes in human RPE, which would have implications in understanding the genetic factors that contribute to AMD.

Using the CRISPRi system, we showed that knockdown of *TMEM97* did not affect cell viability of ARPE19 under normal physiological environment. On the other hand, in the presence of oxidative stress, *TMEM97* knockdown provided protective effects against cell death in ARPE19. Accumulation of oxidative stress is known to play a key role in AMD pathogenesis and progression, which can be caused by multiple risk factors including aging, light damage and cigarette smoking [23]. Interestingly, a recent transcriptome-wide association study using data from 27 human tissues predicted a higher expression level of *TMEM97* in AMD cases compared with controls [24]. This finding is consistent with our results that reduced levels of *TMEM97* have a protective effect on oxidative stress-induced cell death in RPE. Together, our results provide the first evidence that *TMEM97* plays an important role in cell survival of human RPE. Future research beyond this study in primary human RPE or induced pluripotent stem cell-derived RPE would confirm the role of *TMEM97* in regulating cell survival. Also, further studies to assess the functional role of *TMEM97* in other retinal cells, such as photoreceptors, would be of high interest. In addition, given the potential for CRISPRi for multiplexed gene silencing [25], it would be interesting to knockdown *TMEM97* together with other AMD-associated genes to determine if these genes have synergistic contribution to retinal degeneration and pathogenesis of AMD.

*TMEM97* was recently identified as the Sigma 2 receptor. Previous studies in cancer cells showed that a number of Sigma 2 receptor ligands induce apoptosis and cytotoxicity by generating reactive oxygen species [8,26]. On the other hand, Sigma 2 receptor antagonists are shown to provide a protective effect following traumatic brain injury [27]. Our study extended these findings and showed that *TMEM97* played a protective role in RPE against oxidative stresses, providing a potential drug target to alleviate RPE degeneration in AMD. In particular, a Sigma 2 receptor antagonist CT1812 has demonstrated a safety profile in the clinic [10] and is currently being tested as a treatment for Alzheimer’s disease in Phase II trial (ClinicalTrials.gov: NCT03507790). Future studies that evaluate the efficacy of Sigma 2 receptor antagonists in RPE could pave the way to development of novel treatments to improve vision in patients with AMD.

In summary, this study presents an *in vitro* human RPE system with integrated CRISPRi that allows gene silencing. We showed that this *in vitro* CRISPRi system can be used for loss-of-function studies of AMD-associated genes and revealed a novel role of *TMEM97* in regulating human RPE viability against oxidative stresses, providing supporting evidence for its role in AMD pathogenesis.

## Acknowledgement

This research was funded by the University of Melbourne (RCBW) and the Centre for Eye Research Australia (RCBW). DU and TN are supported by the Melbourne Research Scholarship. The Centre for Eye Research Australia receives operational infrastructure support from the Victorian Government.

---

**Supplementary figure 1:**
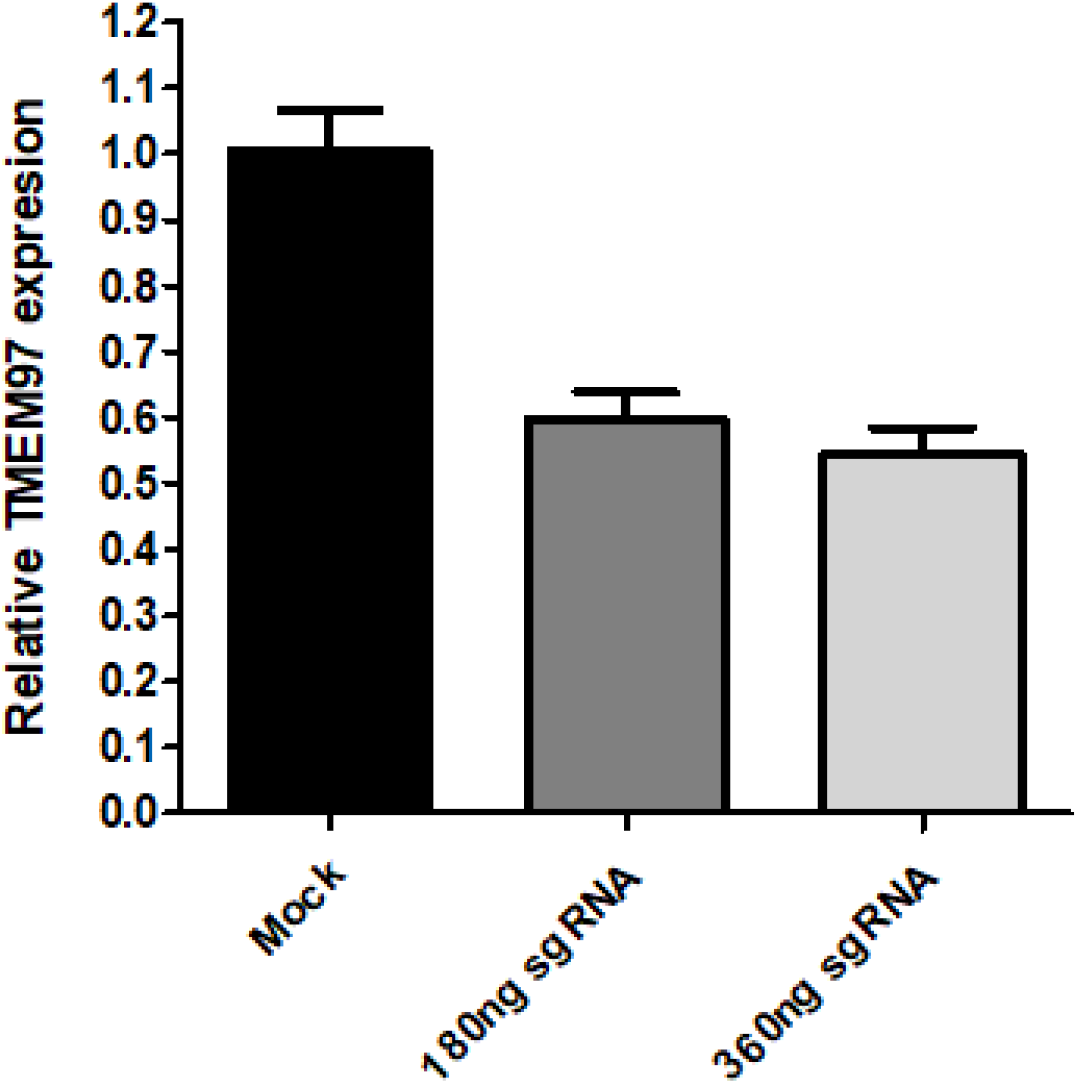
qPCR analysis of TMEM97 expression in ARPE19-KRAB transfected with two different doses of sgRNA (180ng or 360ng sgRNA/well in a 12 well plate). Results are presented as the mean ± SEM of three technical repeats.

**Supplementary figure 2:**
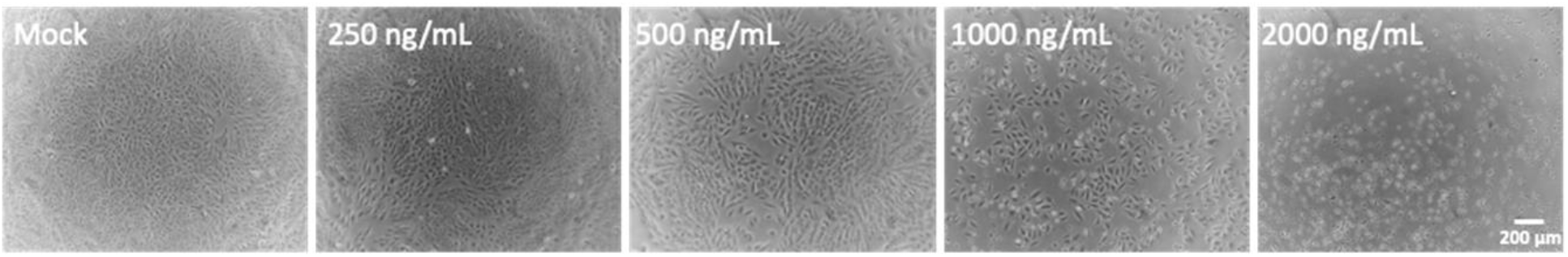
Puromycin dosage study in ARPE19 cells. Representative pictures of ARPE19 treated with various doses of puromycin treatment after 48 hours. Scale bar = 200μm.

**Supplementary table 1:** Details for donor RPE samples used for RNAseq.

| Retina | Donor ID | Eye bulb | Age (yrs) | Sex | Retrieval time (hrs) | Ocular history |
| --- | --- | --- | --- | --- | --- | --- |
| 1 | 17-080 | Right | 77 | F | 4 hrs | Healthy |
| 2 | 17-172 | Left | 69 | F | 4 hrs | Healthy |
| 3 | 18-011 | Left | 71 | M | 6 hrs | Bilateral intraocular lens, exophthalmos |

**Supplementary table 2:**
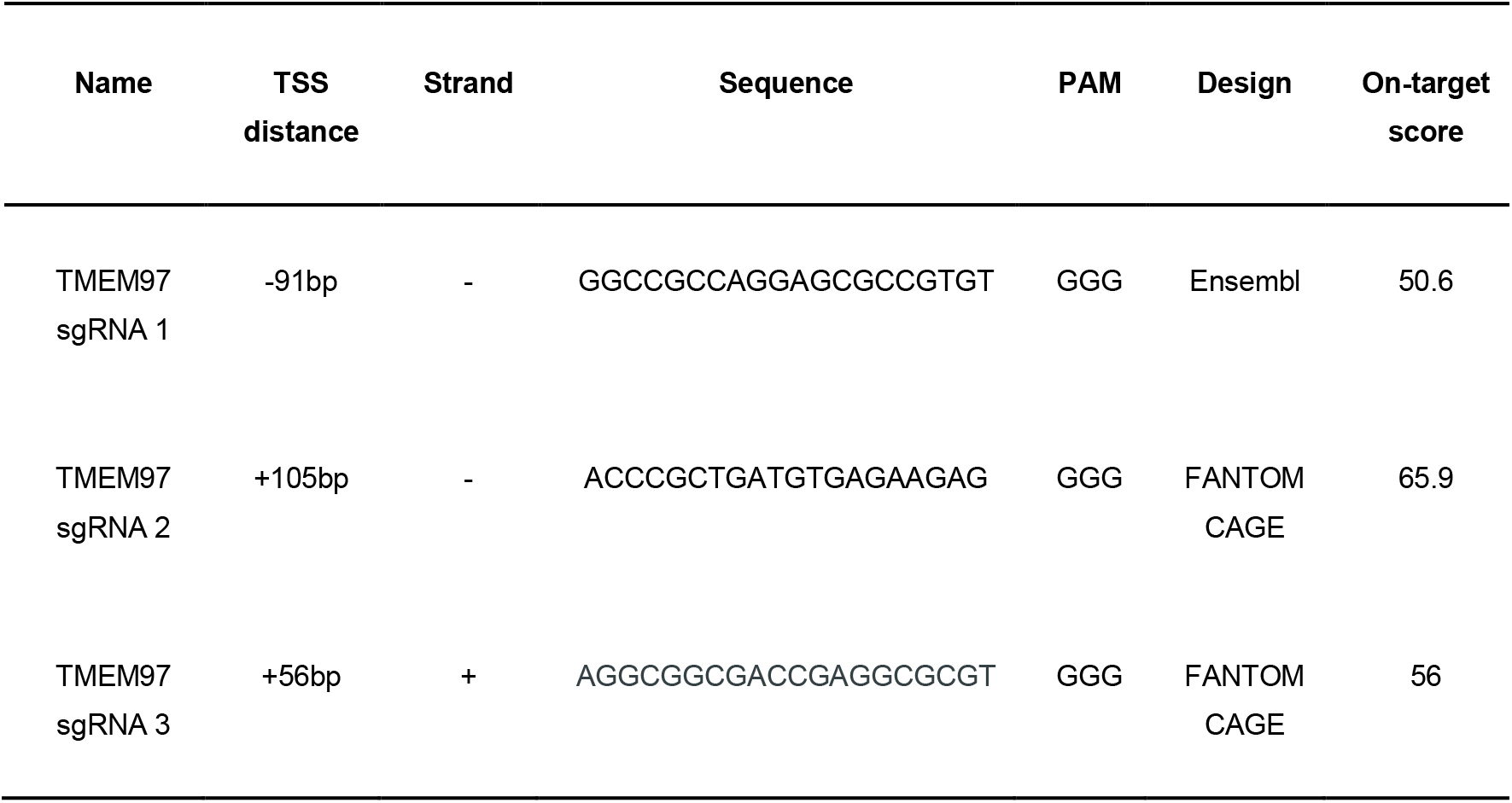
Information of sgRNAs used in this study. TSS distance is based on the transcription start site defined by Ensembl. On-target score is based on [28].

## References

[1] Wong WL, Su X, Li X, Cheung CMG, Klein R, Cheng C-Y, et al. Global prevalence of age-related macular degeneration and disease burden projection for 2020 and 2040: a systematic review and meta-analysis. Lancet Glob Health 2014;2:e106–16.

[2] Fritsche LG, Igl W, Bailey JNC, Grassmann F, Sengupta S, Bragg-Gresham JL, et al. A large genome-wide association study of age-related macular degeneration highlights contributions of rare and common variants. Nat Genet 2016;48:134–43.

[3] Neale BM, Fagerness J, Reynolds R, Sobrin L, Parker M, Raychaudhuri S, et al. Genome-wide association study of advanced age-related macular degeneration identifies a role of the hepatic lipase gene (LIPC). Proc Natl Acad Sci U S A 2010;107:7395–400.

[4] Logue MW, Schu M, Vardarajan BN, Farrell J, Lunetta KL, Jun G, et al. Search for age-related macular degeneration risk variants in Alzheimer disease genes and pathways. Neurobiol Aging 2014;35:1510.e7–18.

[5] Yu Y, Bhangale TR, Fagerness J, Ripke S, Thorleifsson G, Tan PL, et al. Common variants near FRK/COL10A1 and VEGFA are associated with advanced age-related macular degeneration. Hum Mol Genet 2011;20:3699–709.

[6] Cheng C-Y, Yamashiro K, Chen LJ, Ahn J, Huang L, Huang L, et al. New loci and coding variants confer risk for age-related macular degeneration in East Asians. Nat Commun 2015;6:6063.

[7] Alon A, Schmidt HR, Wood MD, Sahn JJ, Martin SF, Kruse AC. Identification of the gene that codes for the σ2 receptor. Proceedings of the National Academy of Sciences 2017;114:7160–5. https://doi.org/10.1073/pnas.1705154114.

[8] Oyer HM, Sanders CM, Kim FJ. Small-Molecule Modulators of Sigma1 and Sigma2/TMEM97 in the Context of Cancer: Foundational Concepts and Emerging Themes. Front Pharmacol 2019;10:1141.

[9] Nicholson HE, Alsharif WF, Comeau AB, Mesangeau C, Intagliata S, Mottinelli M, et al. Divergent Cytotoxic and Metabolically Stimulative Functions of Sigma-2 Receptors: Structure-Activity Relationships of 6-Acetyl-3-(4-(4-(4-fluorophenyl)piperazin-1-yl)butyl)benzo[d]oxazol-2(3H)-one (SN79) Derivatives. Journal of Pharmacology and Experimental Therapeutics 2019;368:272–81. https://doi.org/10.1124/jpet.118.253484.

[10] Grundman M, Morgan R, Lickliter JD, Schneider LS, DeKosky S, Izzo NJ, et al. A phase 1 clinical trial of the sigma-2 receptor complex allosteric antagonist CT1812, a novel therapeutic candidate for Alzheimer’s disease. Alzheimers Dement 2019;5:20–6.

[11] Davidson M, Saoud J, Staner C, Noel N, Luthringer E, Werner S, et al. Efficacy and Safety of MIN-101: A 12-Week Randomized, Double-Blind, Placebo-Controlled Trial of a New Drug in Development for the Treatment of Negative Symptoms in Schizophrenia. Am J Psychiatry 2017;174:1195–202.

[12] Hung SSC, McCaughey T, Swann O, Pébay A, Hewitt AW. Genome engineering in ophthalmology: Application of CRISPR/Cas to the treatment of eye disease. Prog Retin Eye Res 2016;53:1–20.

[13] Dominguez AA, Lim WA, Qi LS. Beyond editing: repurposing CRISPR-Cas9 for precision genome regulation and interrogation. Nat Rev Mol Cell Biol 2016; 17:5–15.

[14] Patro R, Duggal G, Love MI, Irizarry RA, Kingsford C. Salmon provides fast and bias-aware quantification of transcript expression. Nat Methods 2017;14:417–9.

[15] Fang L, Hung SSC, Yek J, El Wazan L, Nguyen T, Khan S, et al. A Simple Cloning-free Method to Efficiently Induce Gene Expression Using CRISPR/Cas9. Mol Ther Nucleic Acids 2019;14:184–91.

[16] Hung SSC, Van Bergen NJ, Jackson S, Liang H, Mackey DA, Hernández D, et al. Study of mitochondrial respiratory defects on reprogramming to human induced pluripotent stem cells. Aging 2016;8:945–57.

[17] Hung SSC, Wong RCB, Sharov AA, Nakatake Y, Yu H, Ko MSH. Repression of global protein synthesis by Eif1a-like genes that are expressed specifically in the two-cell embryos and the transient Zscan4-positive state of embryonic stem cells. DNA Res 2013;20:391–402.

[18] Lukowski SW, Lo CY, Sharov AA, Nguyen Q, Fang L, Hung SS, et al. A single-cell transcriptome atlas of the adult human retina. EMBO J 2019;38:e100811.

[19] Gilbert LA, Larson MH, Morsut L, Liu Z, Brar GA, Torres SE, et al. CRISPR-mediated modular RNA-guided regulation of transcription in eukaryotes. Cell 2013;154:442–51.

[20] Radzisheuskaya A, Shlyueva D, Müller I, Helin K. Optimizing sgRNA position markedly improves the efficiency of CRISPR/dCas9-mediated transcriptional repression. Nucleic Acids Res 2016;44:e141.

[21] Datta S, Cano M, Ebrahimi K, Wang L, Handa JT. The impact of oxidative stress and inflammation on RPE degeneration in non-neovascular AMD. Prog Retin Eye Res 2017;60:201–18.

[22] Bringmann A, Syrbe S, Görner K, Kacza J, Francke M, Wiedemann P, et al. The primate fovea: Structure, function and development. Prog Retin Eye Res 2018;66:49–84.

[23] Bellezza I. Oxidative Stress in Age-Related Macular Degeneration: Nrf2 as Therapeutic Target. Front Pharmacol 2018;9:1280.

[24] Strunz T, Lauwen S, Kiel C, International AMD Genomics Consortium (IAMDGC), den Hollander A, Weber BHF. A transcriptome-wide association study based on 27 tissues identifies 106 genes potentially relevant for disease pathology in age-related macular degeneration. Sci Rep 2020;10. https://doi.org/10.1038/s41598-020-58510-9.

[25] McCarty NS, Graham AE, Studená L, Ledesma-Amaro R. Multiplexed CRISPR technologies for gene editing and transcriptional regulation. Nat Commun 2020;11:1281.

[26] Liu C-C, Yu C-F, Wang S-C, Li H-Y, Lin C-M, Wang H-H, et al. Sigma-2 receptor/TMEM97 agonist PB221 as an alternative drug for brain tumor. BMC Cancer 2019;19:473.

[27] Vázquez-Rosa E, Watson MR, Sahn JJ, Hodges TR, Schroeder RE, Cintrón-Pérez CJ, et al. Neuroprotective Efficacy of a Sigma 2 Receptor/TMEM97 Modulator (DKR-1677) after Traumatic Brain Injury. ACS Chem Neurosci 2019;10:1595–602.

[28] Doench JG, Fusi N, Sullender M, Hegde M, Vaimberg EW, Donovan KF, et al. Optimized sgRNA design to maximize activity and minimize off-target effects of CRISPR-Cas9. Nat Biotechnol 2016;34:184–91.

